# Neuromolecular responses in disrupted mutualistic cleaning interactions under future environmental conditions

**DOI:** 10.1101/2023.07.20.549851

**Authors:** S. Ramírez-Calero, J. R. Paula, E. Otjacques, T. Ravasi, R. Rosa, C. Schunter

**Author notes:** Correspondence: Celia Schunter.

## Abstract

Mutualistic interactions, which constitute some of the most advantageous interactions among fish species, are highly vulnerable to environmental changes. A key mutualistic interaction is the cleaning service rendered by the cleaner wrasse, *Labroides dimidiatus*, which involves intricate processes of social behaviour to remove ectoparasites from client fish and can be altered in near-future environmental conditions. Here, we evaluated the neuromolecular mechanisms behind the behavioural disruption of cleaning interactions in response to future environments. We subjected cleaner wrasses and surgeonfish (*Acanthurus leucosternon*, serving as clients) to elevated temperature (Warming, 32°C), increased levels of CO_2_ (High CO_2_, 1000 ppm), and a combined condition of elevated CO_2_ and temperature (Warming & High CO_2_, 32°C & 1000 ppm) for 28 days. Each of these conditions resulted in behavioural disruptions concerning the motivation to interact and the quality of interaction (High CO_2_ -80.7%, Warming – 92.6%, Warming & High CO_2_ -79.5%, *p*<0.001). Using transcriptomic of the fore-, mid-, and hindbrain, we discovered that most transcriptional reprogramming in both species under warming conditions occurred primarily in the hind- and forebrain. The associated functions under warming were linked to stress, heat shock proteins, hypoxia, and behaviour. In contrast, elevated CO_2_ exposure affected a range of functions associated with GABA, behaviour, visual perception, and circadian rhythm. Interestingly, in the combined Warming & High CO_2_ condition, we did not observe any expression changes of behaviour. However, we did find signs of endoplasmic reticulum stress and apoptosis, suggesting not only an additive effect of the environmental conditions but also a trade-off between physiological performance and behaviour in the cleaner wrasse. We suggest that impending environmental shifts can affect the behaviour and molecular processes that sustain mutualistic interactions between *L. dimidiatus* and its clients, which could have a cascading effect on their adaptation potential and possibly cause large-scale impacts on coral reef ecosystems.

## INTRODUCTION

Global change is one of the main drivers of marine biodiversity decline (Nagelkerken & Munday, 2016). Rapid modifications of temperature and pH in the oceans are accelerating demographic changes in marine species, substantially modifying ecosystem diversity, and affecting interspecific interactions (Burrows et al., 2019; Nagelkerken et al., 2018). In particular, key ecological interactions such as competition, reproduction and mutualism are modified by environmental conditions that consequently generate new social contexts among species (Nagelkerken & Munday, 2016). Therefore, there is a strong link between the physical environment and the outcome of ecological processes mediated by animal behaviour. Also, the effect of environmental changes on marine organisms has potentially profound outcomes at higher levels of biological organization (Nagelkerken & Munday, 2016).

Changes in the marine environment such as elevated levels of CO_2_ (ocean acidification) and elevated temperature (ocean warming) have a variety of effects on marine organisms, especially in vertebrates such as fish (Clements & Hunt, 2015; Milazzo et al., 2013; Rosa et al., 2017; Santos et al., 2021). CO_2_ changes in the water lead to acid-base regulation in animal tissues influencing ecological traits such as reproduction, larval growth, and biological senses that can have consequences on fish boldness, learning, recognition, and predator avoidance, among others (Nagelkerken & Munday, 2016). For instance, levels of monoamines, such as dopamine and serotonin, can be chemically altered through ion exchanges, and have been shown to contribute to fish aggression and decision-making (Demin et al., 2019; Paula et al., 2019a). However, there are inter-specific and intra-specific variations in the sensitivity to ocean acidification which may allow for adaptive physiological strategies in response to an acidified ocean (Kang et al., 2022). At present, behavioural impairment through elevated CO_2_ has mostly been attributed to the alteration in Gamma Aminobutyric Acid (GABA) neurotransmission in the brain (Munday et al., 2019; Schunter et al., 2019). However, additional mechanisms such as the alteration of Ca^2+^/calmodulin protein kinase II (CaMKII) affecting olfactory abilities and the movement of AMPA receptors in neurons to generate synaptic plasticity have also been explored (Porteus et al., 2018).

Further effects of global change, such as elevated temperature, generate changes in the metabolic rates of fish, reducing their aerobic performance and collapsing aerobic capabilities that are required for basic functions such as obtaining food or maintaining symbiosis with other species (Milazzo et al., 2013; Payne et al., 2016). Since there is a physiological need to cope with warming conditions to avoid heat shocks, proteins and transcription factors can be activated to protect and maintain cellular functions (Bernal et al., 2020). In fact, the expression of cellular stress responses with genes associated with antioxidant defense, apoptosis, and protein folding are often exhibited with an elevated temperature (Somero, 2020), and the effect of this elevated temperature response is a reduction in fish performance across immune responses, respiration and foraging (LeBlanc et al., 2011; Pörtner & Farrell, 2008; Rendell et al., 2006; Samaras et al., 2018; Tomalty et al., 2015). Furthermore, high temperatures also interact with elevated CO_2_ and increase, reduce or change the effects on metabolic demands, growth, development, behaviour and survival of marine fish species (Allan et al., 2017; Biro et al., 2010; Pörtner & Farrell, 2008; Rosa et al., 2014). For example, with the combination of environmental changes, antipredator behaviour is reduced in damselfishes (Ferrari et al., 2015). On the other hand, anemonefish food consumption increases under exposure to both temperature and CO_2_, whereas no effects are seen with CO_2_ exposure alone (Nowicki et al., 2012).

Cleaning mutualisms, one of the most beneficial interactions between fish species, is highly susceptible to environmental changes (Paula et al., 2019a; Paula, et al., 2019b; Paula et al., 2022; Paula et al., 2023; Triki et al., 2018). In particular, cleaning mutualisms are a crucial interaction on coral reef ecosystems consisting of the removal of ectoparasites and dead tissue from the skin of other fishes (known as ‘clients’), enhancing their health and survival (Bshary & Côté, 2008; Grutter, 1999). For instance, the bluestreak cleaner wrasse *Labroides dimidiatus* is known for its cleaning ability that leads to the enhancement of reef fish well-being and diversity (Waldie et al., 2011). This species also possesses remarkable cognitive abilities of learning and memory (Masanori et al., 2018) which allows the establishment of long-term mutualistic relationships with clients that visit cleaning stations to obtain stress relief (Soares et al., 2011) and boost their health (Ros et al., 2011), while the cleaner wrasse obtains food in return. Therefore, changes to the interaction behaviour of *L. dimidiatus* due to environmental conditions could have consequences for marine fish communities (Grutter, 1999; Waldie et al., 2011). Disruptions such as habitat degradation and temperature changes disturb mutualistic interactions leading to shifts from mutualism to antagonism, loss of interaction, and unexpected switches to new participants or partners (Kiers et al., 2010). For instance, ocean acidification and warming impair the motivation to interact of cleaner wrasses, indirectly leading to cascading effects in fish communities and the abundance of clients (Paula et al., 2019a; Waldie et al., 2011). In addition, the quality of these interactions can be affected by the interruption of cognitive processes involved in recognition of individual clients, which is essential in establishing cleaning relationships (Tebbich et al., 2002). Consequently, this may affect the functioning of the mesolimbic reward system (Soares, 2017a) and lead to mutualism breakdown which indirectly impact coral reef ecosystems by decreasing reef fish diversity (Waldie et al., 2011).

Previously, Paula et al. (2019) found that when cleaner wrasses were exposed to warmer temperatures and high CO_2_ levels for 45 days, they interacted less frequently with surgeonfish clients. Furthermore, the cleaners used more reconciliation strategies, such as providing tactile stimulation, without increasing any dishonest behaviour (*i.e.* ‘cheating’, behaviour known as cleaner’s preference to eat mucus instead of cleaning ectoparasites (Grutter & Bshary, 2003). This suggested that the cleaners were no longer able to anticipate the costs of interacting with the clients. The behavioural disruptions in cleaning interactions are known to be correlated with changes in the levels of dopamine and serotonin in multiple brain regions (Paula et al., 2019b) and to be reverted by the administration of GABAergic antagonists (Paula et al., 2023). However, to clearly understand the mechanisms behind these disruptions, a molecular approach is needed.

To systematically study these underlying mechanistic changes in interacting cleaner wrasse *L. dimidiatus* and its client (*Acanthurus leucosternon*) we observed the interaction behaviour for fish exposed to i) present-day environmental ‘control’ conditions (29 °C, pCO_2_ ∼400 ppm), ii) ‘Warming’ (32 °C, pCO_2_ ∼400 ppm), iii) ocean acidification ‘High CO_2_’ (29 °C, pCO_2_ ∼1000 ppm) or iv) elevated ‘Warming & High CO_2_’ (32 °C, pCO_2_ ∼1000 ppm), following IPCC’s RCP scenario 8.5 (figure 1, table S1). We evaluated the fine-scale transcriptional responses across the three main regions of the brain (fore-, mid- and hindbrain) known to harbour the expression of significant neurotransmission, neurohormones and neuropeptides during cleaning interactions (Paula et al., 2019b; Soares et al., 2010). Since cleaning interactions involve the expression of the dopaminergic and glutamatergic pathways (Ramírez-Calero et al., 2021), we may expect brain molecular drivers in the response to environmental change in the cleaner wrasse and its client to include cellular stress response, changes to GABAergic neurotransmission and in gene expression levels of neuroamines. Due to the importance of this interspecific and mutualistic interaction, it is necessary to unravel the underlying mechanisms that drive such interactions. Moreover, it is also essential to understand the mechanisms that cause a disruption to these behaviours, which potentially result in mutualism breakdown and wide-ranging effects on the coral reef ecosystem.

**Figure 1.**
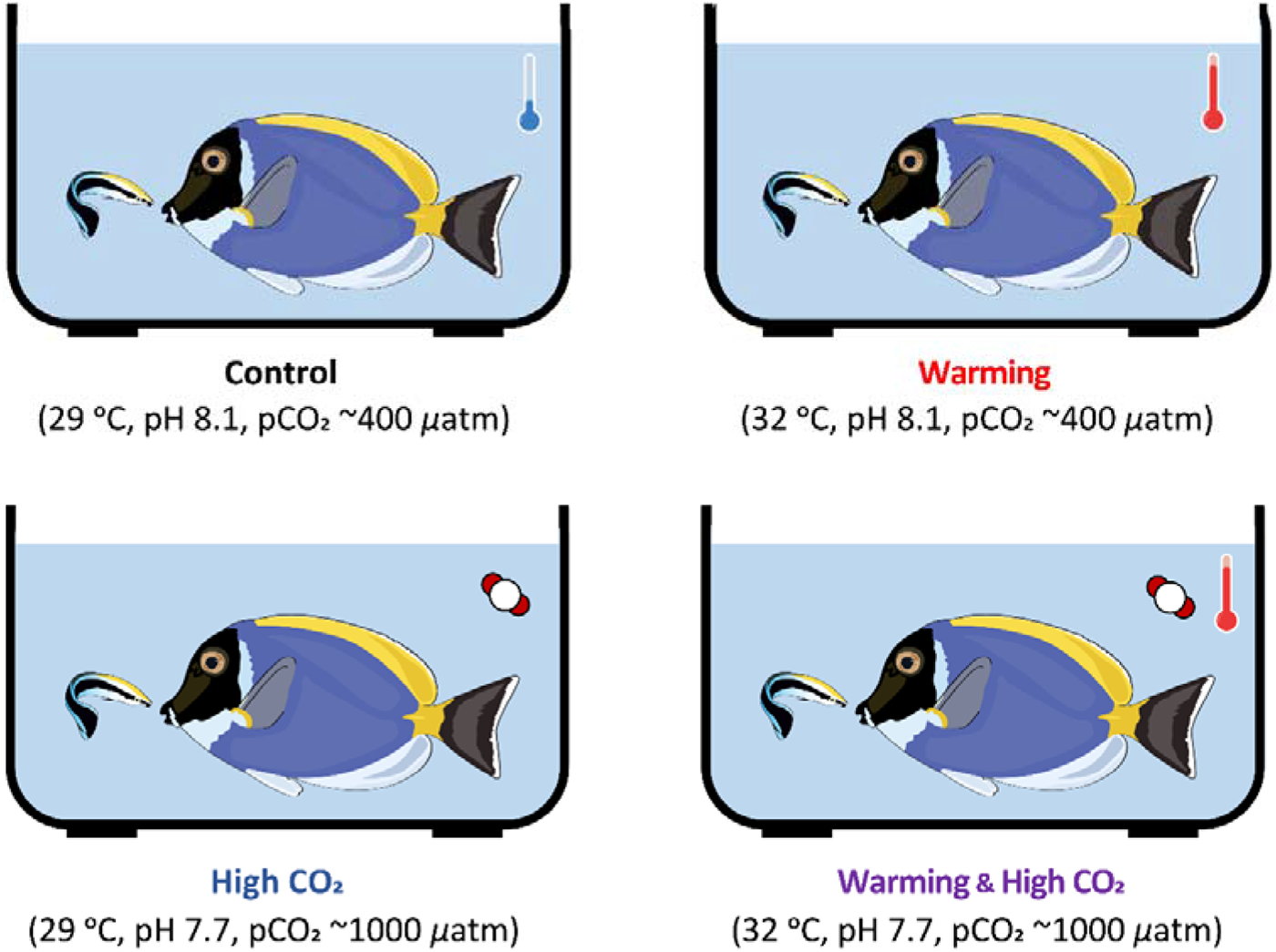
Experimental design where *Labroides dimidiatus* (N=24) and *Acanthurus leucosternon* (N=24) were allowed to interact after an environmental acclimation period (28 days) in one of the following conditions: Control, Warming, High CO_2_, or a combined condition of Warming & High CO_2_ (aquarium setup parameters and behavioural data can be found on table S2-S3a, b).

## RESULTS

### Behavioural responses

After 28 days of acclimation to one of the different treatment scenarios, we note that the proportion of time spent interacting was significantly affected by the interaction of CO_2_ and Temperature (*p*=0.002; table S3b). *Post hoc* comparisons further revealed that time spent in cleaning interactions decreased significantly with High CO_2_ (-66.5%, *p=*0.01) and Warming (-75.5%, *p=*0.01), but not for Warming & High CO_2_ (*p*=0.112; figure 2a, table S3c). Considering cleaners’ motivation to interact, the proportion of interactions started by cleaners was also significantly affected by the interaction of CO_2_ and Temperature (*p*<0.001; table S3b). *Post hoc* comparisons indicated a significant decrease under all treatments compared to the control (High CO_2_ -80.7%, *p*<0.001; Warming -92.6%, *p*<0.001; Warming & High CO_2_ -79.5%, *p*<0.001; figure 2b, table S3c). Contrarily, clients’ motivation (client posing displays ratio, figure 2c) was significantly altered by Temperature (*p*<0.001) but not CO_2_ (*p*=0.771) nor the interaction of CO_2_ and temperature (*p*=0.187, table S3b). *Post hoc* comparisons (table S3c) revealed that treatments with high temperature had significantly higher client posing displays ratio than under control temperatures (Warming +246.2%, *p*=0.002; Warming & High CO_2_ + 198.7% *p*=0.006), but not under high CO_2_ only (*p*=0.268).

When considering the quality of the cleaning interactions, namely the proportion of interaction time spent in tactile stimulation, there was no significant change attributable to temperature, CO_2_, or the interaction between CO_2_ and temperature (figure 2d, table 3c). Additionally, while the interaction between CO_2_ and temperature significantly influenced the ratio of client jolts per 100 seconds of interaction, subsequent *post hoc* analysis revealed that none of the treatment conditions showed a significant deviation from the control (figure 2e, table 3c. On the contrary, the mean interaction duration experienced a significant reduction in response to the combined effects of CO_2_ and temperature (figure 2f, *p*=0.016). Further *post hoc* comparisons revealed a marked decrease in interaction duration under all experimental conditions compared to the control group (table S3c). Specifically, the High CO_2_ condition led to a decrease of 57.7% (*p*=0.007), the Warming condition to a reduction of 72.6% (*p*=0.002), and the combination of both Warming & High CO_2_ resulted in a 66.1% decrease (*p*=0.002).

**Figure 2.**
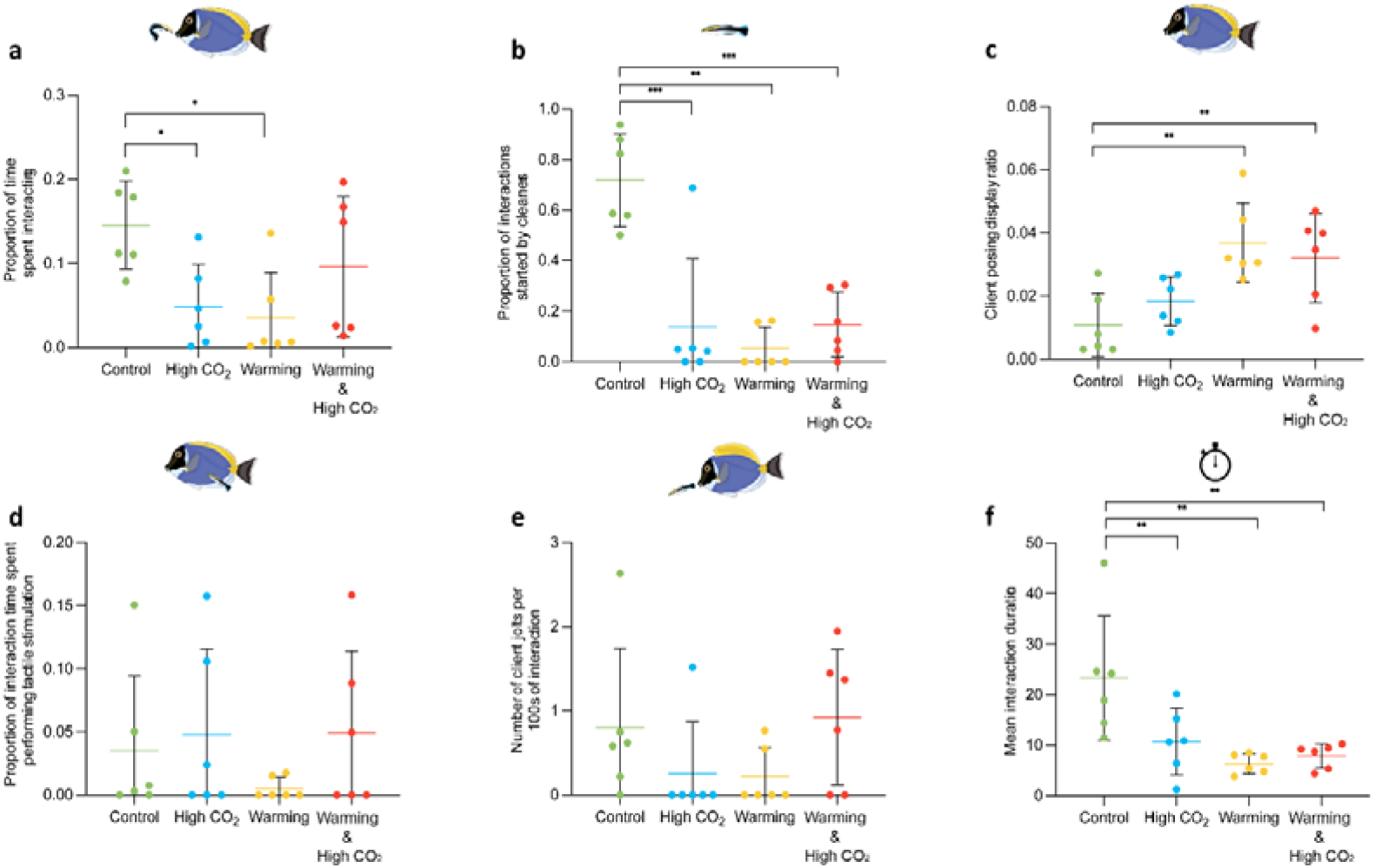
Behavioural responses from interaction trials between the cleaner *L. dimidiatus* and client *A. leucosternon*. a) Proportion of time interacting (in seconds), b) Proportion of interactions started by cleaners, c) Client posing display ratio, d) Proportion of time spent in tactile stimulation (in seconds), e) Number of client jolts per 100s interaction, f) Mean interaction duration (in seconds). (*) define significance based on *post hoc* tests (*p*<0.05). Additional details of associated tests are found in table S3b-c.

### Transcriptional response

Whole transcriptional response (number of differentially expressed genes – DEGs) following the interaction trial revealed a similar pattern in both species across treatments: Warming > Warming & High CO_2_ > High CO_2_. Warming showed the largest molecular response for *L. dimidiatus* (5,804 DEGs) and *A. leucosternon* (3,493 DEGs) (figure 3), followed by Warming & High CO_2_ with 4,581 DEGs for *L. dimidiatus* and 505 DEGs for *A. leucosternon.* Finally, High CO_2_ alone showed the smallest numbers of DEGs for both species revealing 2,508 for *L. dimidiatus* and 376 for *A. leucosternon* (figure 3). For *L. dimidiatus*, the 672 DEGs common across all environmental conditions (regardless of brain region) are involved in protein folding processes, positive regulation of endothelial cell proliferation, chemosensory behaviour, regulation of bone mineralization, among others (table S4, figure 3a). As for *A. leucosternon*, the 30 common DEGs were related with biological regulation, regulation of cellular process and regulation of apoptotic signalling pathway (table S5, figure 3b).

**Figure 3.**
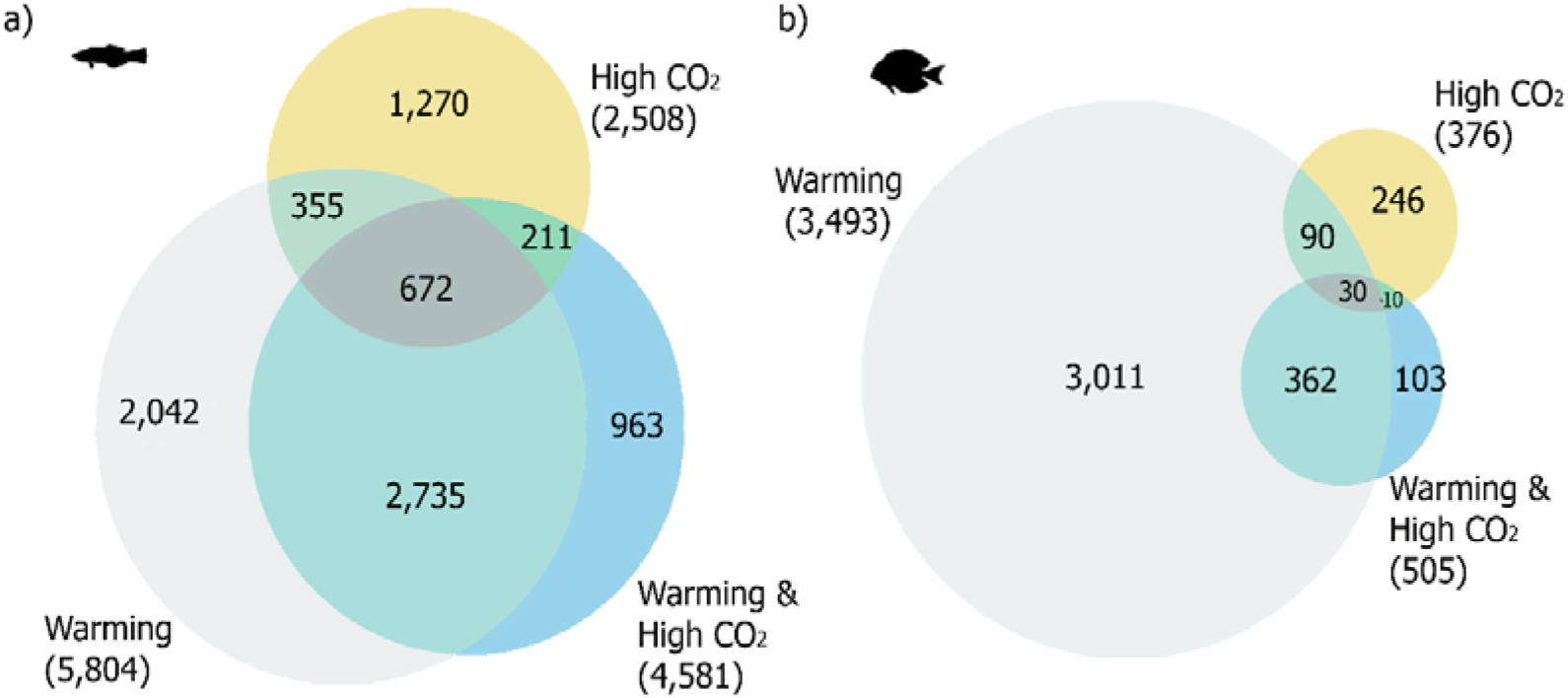
Number of differentially expressed genes that are unique to each condition and common between elevated temperature, High CO_2_ and elevated Warming & High CO_2_ conditions of the whole brain for (a) the cleaner fish *Labroides dimidiatus* and (b) the powder-blue surgeonfish *Acanthurus leucosternon*. Numbers in brackets are proportional to the size of the circle and represent the total differential expressed genes found under each environmental condition.

Even though the magnitude of transcriptional reprogramming in the whole brain associated with the conditions was similar for both studied species, differences in gene expression patterns were exhibited across brain regions (figure 4, figure S1-S3). In particular, for *L. dimidiatus*, the hindbrain region (HB) displayed 1,198 unique DEGs for High CO_2_, while in *A. leucosternon* more DEGs were found in the forebrain (204, FB) under the same conditions. Furthermore, the FB region in *L. dimidiatus* presented the largest molecular reaction under the Warming condition (1,632), while in *A. leucosternon* was in the HB (2,732). Finally, under Warming & High CO_2_, most differential expression in *L. dimidiatus* was observed in FB region (471), while in *A. leucosternon* was in HB (82) (figure 4a-b, figures S1-S6).

**Figure 4.**
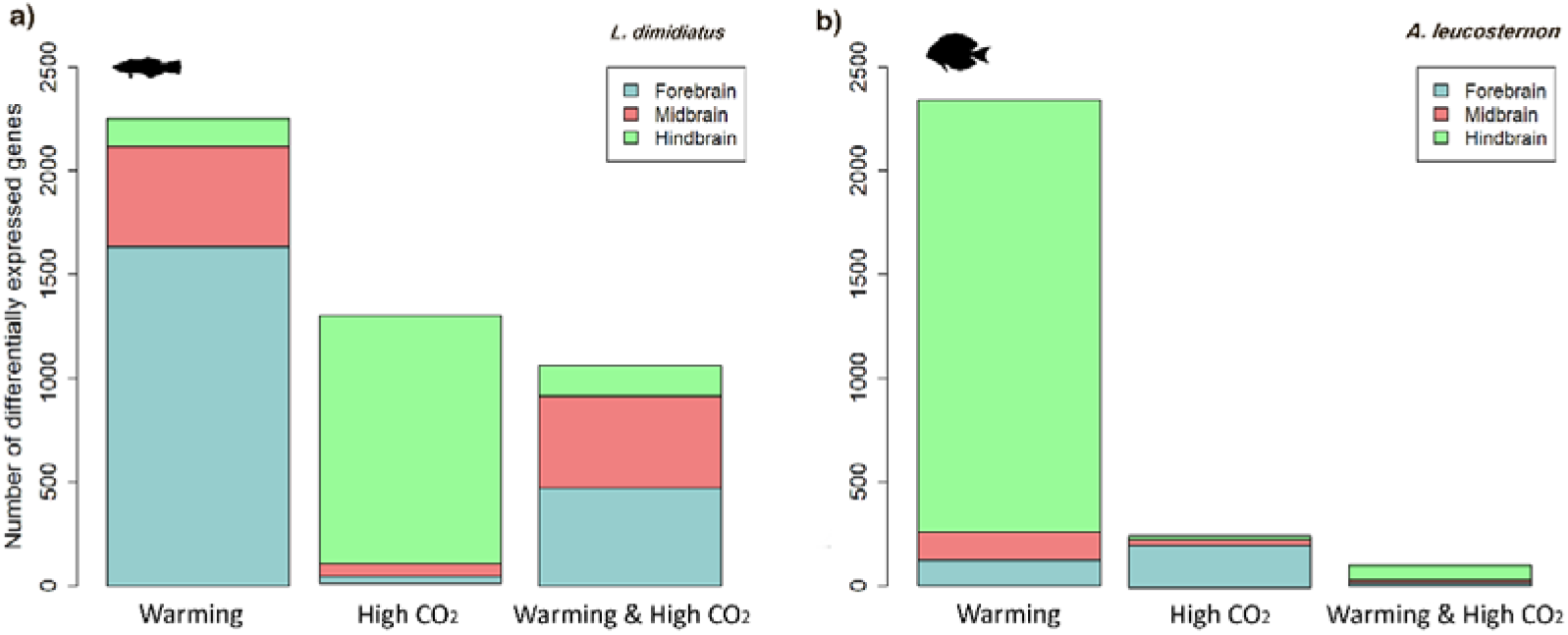
Number of unique differentially expressed genes (DEGs) for each brain region under each environmental condition compared to the control condition for (a) *L. dimidiatus* and (b) *A. leucosternon*. Further overlapping DEGs between brain regions can be found in figure S1-S6.

### Molecular responses to elevated temperature

#### a) Labroides dimidiatus

Exposure to a higher temperature led to unique functional changes (only seen for this treatment). These changes were related to immune response, execution phase of apoptosis, response to hypoxia, and stress-activated MAPK cascade (table S6a). Furthermore, elevated temperature elicited specific changes in FB (1,450 DEGs) and MB (309 DEGs) (figure 4a, Table S6b-c), characterized by functional enrichments of cellular stress responses and the expression of actin filaments genes, DNA damage, glucose, and glycine receptors, response to environmental stress, cell proliferation and heat shock proteins (table S6a-c). In particular, genes underlying functions of stress and hypoxia were evidenced by the upregulation of HSF2 (*Heat shock factor protein 2*), a specific promoter under conditions of heat stress, processes of insulin metabolism given by IGF1R (*insulin growth factor 1*), AKT3 (*RAC-gamma serine/threonine-protein kinase*) and INSR (*insulin receptor*) and transcriptional co-suppressor functions of hypoxia through HIPK2 (*Homeodomain-Interacting Protein Kinase 2*). Regarding the HB specifically, we found receptors for the perception of pain differentially expressed, such as OPRK (O*pioid Receptor Kappa* 1). Simultaneously with these stress signatures, histones, and epigenetic regulation were also differentially expressed under Warming. Peptide hormone secretion and thyroid hormone receptor binding were upregulated mainly in the FB, underlined by upregulation of hormone receptors to behavioural responses to stress (*Corticotropin Releasing Hormone Receptor 2*) and adenylate cyclase (cAMP) inhibition (*Cannabinoid Receptor 1*, also Differentially Expressed in MB). Furthermore, neurotransmission was differentially regulated with genes related to processes of calcium (CAC1D, KCC2B, SORCN), glutamate (DHE3), GABA (GABR1), and potassium (KCNB1) transport. Finally, molecular signatures of olfactory behaviour, adult locomotory behaviour, and social behaviour were evident almost exclusively in FB (table S6a-c).

#### b) Acanthurus leucosternon

For the client species, stress responses, metabolic functions, and synapse activity were also elicited with elevated temperature (3,493 DEGs; table S7, figure 3b). Although, in contrast to the cleaner wrasse (figure 4a), differential gene expression in HB was the largest among the brain regions for *A. leucosternon* (2,732, figure 4b), and similar functional processes were shared with FB. For instance, responses to stimulus, epigenetic regulation, synapse activity, behaviour, and learning were differentially expressed for the client species under Warming. On the other hand, some DEGs were common with the cleaner wrasse, such as adult locomotory behaviour, locomotory exploration behaviour and social behaviour. However, their differential expression was significant almost exclusively in the HB (table S7). Additionally, the FB and MB revealed changes in molecular signatures involved in synaptic transmission (glutamate and GABA), protein binding, and cellular responses to stimuli (table S7) and highlighted a significant alteration of metabolic process and glutamatergic synapses under elevated temperature. Some enriched hormone responses were found related to thyroid hormone receptor binding, regulation of growth hormone secretion, and steroid hormone binding, underlined by upregulation of *Thyroid hormone receptor alpha* (THRA) and several *Chromodomain-helicase-DNA-binding proteins* (CHD6, 7, 8, 9), found almost exclusively in the HB (table S7).

### Molecular responses to High CO_2_ exposure

#### a) Labroides dimidiatus

Under High CO_2_, the differentially expressed molecular signatures in *L. dimidiatus* also showed to initiate cellular stress responses. The molecular signatures displayed differed from the other treatments, with the HB presenting the largest number of DEGs compared to the other brain regions under High CO_2_ (HB: 1,198 > MB: 60 > FB: 32, figure S2). The functional changes were related to stress, such as positive regulation of stress-activated MAPK cascade and response to osmotic stress in the HB (table S8). Molecular responses in apoptosis, osmotic and oxidative stress were also found in High CO_2_. However, the DEGs involved differed from those in Warming (table S8, figure 5). Unique functional responses to High CO_2_ were related to synaptic neurotransmission, glutamate and GABA such as positive regulation of synaptic transmission, glutamatergic and gamma-aminobutyric acid secretion. These processes were underlined by several upregulated ionotropic (NMDE1, NMDZ1) and downregulated metabotropic glutamate receptors (GRM8, 2), Gamma-aminobutyric acid associated-genes (GBRAP, GABT) and Sodium-chloride transporters (SC6A1, 13). Further processes related to circadian rhythm and visual perception were found (e.g. camera-type eye photoreceptor cell differentiation, eye photoreceptor cell fate commitment, and blue light photoreceptor activity). These processes were differentially expressed mainly in the HB, with a downregulation of phosphodiesterases (PLCB1) and phosphatases (PHR4B) as well as an upregulation of cristallins (CRYAB), cryptochromes (CRY1, 2), sidekick (SDK2) and calcium proteins (CABP1). Furthermore, hormone responses (e.g. dopamine, gonadotropin, estrogen, thyroid and corticotropin; table S8) were also found with functional enrichments in the HB and upregulated DEGs of corticotropin receptor and releasing factors (CRF, CRHBP), dopamine genes (DRD5, TY3H) and Melatonin receptors (MR1BB). Finally, epigenetic regulations were found, such as histone regulation, acetylation and methylation processes (table S8), as well as a variety of enriched behaviours, learning and memory functions (e.g. adult locomotory behaviour, social behaviour, olfactory and vocal learning, short-, medium- and long-term memory). These functions were underlined by upregulated transcription in the HB of glutamate receptors (GRIA2, 3, 4, NMDZ1), dopamine genes (TY3H, DRD5), Isotocin receptor (ITR), and Early growth response protein 1 (EGR1, table S8).

**Figure 5.**
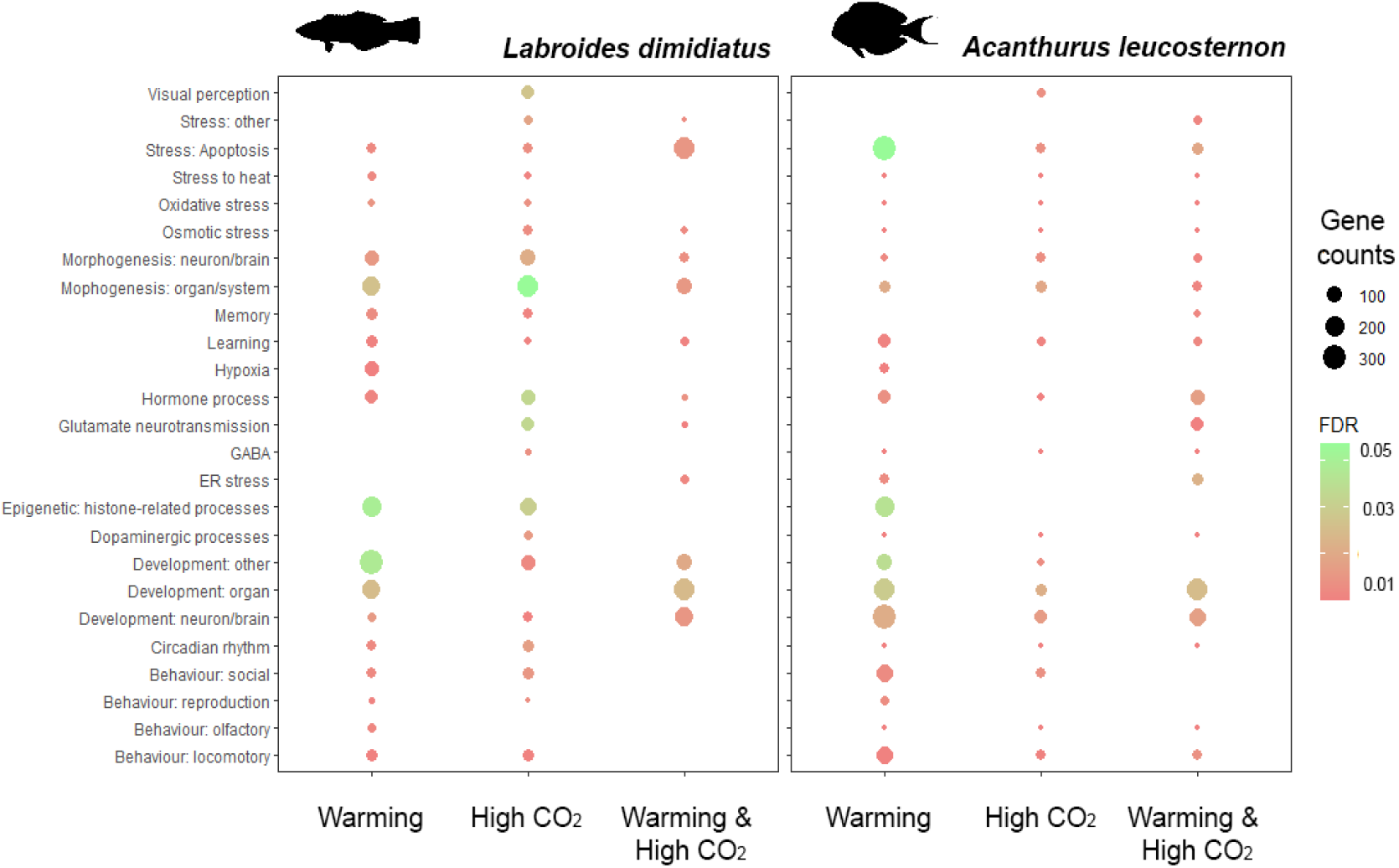
Significantly enriched functions in across brain regions of *L. dimidiatus* and *A. leucosternon* with different near-future environmental treatments (Warming, High CO_2,_ and Warming & High CO_2_). The size of the circles is proportional to the number of differentially expressed genes, and the colour of the circles represents the false discovery rate adjusted for enrichment significance level (FDR <0.05).

#### b) Acanthurus leucosternon

High CO_2_ generated the most variable effect in the molecular reaction of the exposed client individuals compared to the other conditions, revealing differential responses among the biological replicates (figure S7). Various genes associated with organ development, stress, diencephalon morphogenesis, behaviour and learning, such as locomotory behaviour and associative learning, were altered in expression, mainly in FB, which presented the largest molecular response (table S9, figure 4b). In addition, MB and HB had fewer unique DEGs than FB (204, figure 4b), and their molecular functions were involved in synapses activation and signalling, peroxidase activity, oxygen carrier activity, gamma-aminobutyric acid signalling pathway, homeostasis and immune responses (table S9). Unique functional responses in High CO_2_ for the client included functions of signalling involved in the determination of organs and tissues, as well as responses to stress. Interestingly, hormone activity was the only enriched function found in this species that had the upregulation exclusively in FB of Isotocin (NEUI), Gonadotropin (GTHB1), Pro-thyrotropin (TRH), Thyrotropin (TSHB) and Somatostatin (SMS1B) hormones. In addition, behavioural processes are altered, including adult locomotory and social behaviour, underlined by upregulated genes of Glutamate decarboxylase (DCE1) and downregulated glutamate receptor GRM5, Isotocin (NEUI) and dopamine-related gene Tyrosine 3-monooxygenase (TY3H), almost exclusively in FB, except for GRM5. Finally, learning processes were altered, but no processes of memory, epigenetic regulation or circadian rhythm were found for this species.

### Molecular response to the combined treatment: Warming & High CO_2_

#### a) Labroides dimidiatus

The combined treatment of Warming & High CO_2_ triggered the differential expression of 4,581 DEGs (figure 3a, table S10a), from which 60% (2,735 DEGs) were shared with Warming treatment. These DEGs were mostly related to associative learning, oxidative stress, cell death, insulin activity, and immune response. DEGs shared with High CO_2_ corresponded to only 4.6% (211 DEGs) and were involved in osmotic stress, long-term synaptic potentiation, and nervous projection development (figure 3a). In Warming & High CO_2_, a total of 963 genes were found to be uniquely differentially expressed (21%, figure 3a), mainly in the FB (471 DEGs) and MB (442 DEGs), and only 146 DEGs in HB (figure S2). In this unique response, elevated stress response signals are revealed by neuron apoptotic process, response to endoplasmic reticulum stress, and regulation of neuron death (table S10b, figure 5). According to the brain regions, FB revealed cellular stress response but also neuron development (table S10b), while genes involved with the regulation of cell population proliferation and regulation of cell death were differentially expressed in MB (table S10b). Concerning HB, genes related to the glutamatergic synapse pathway, calcium/calmodulin-dependent serine protein kinase were differentially expressed (table S10). Furthermore, Warming & High CO_2_ unique responses were related to organ development and morphogenesis and the genes were mostly transcribed in FB and MB, underlined by upregulation of DNA-binding transcription of helix-loop-helix processes (ID3A), cysteine glutamate transporters (XCT) and receptors (GRID1, NMDE4, GRM3, 8). Furthermore, there was an upregulation of organ and neuromuscular tissues development (BMR1B, HDAC8, FGF12, TSN2, FZD2-6, AGRIN), synaptic integrity and signalling (GRB2A, PTPRF, CSKP, CBLN1), as well as histone-related genes (KDM1B, HDAC8). In addition, steroid and phosphatidic acid processes with upregulations of glutamate transporter XCT and downregulation of glutamate receptor GRM3 and insulin (INSI1) were observed, but also by steroid hormone receptor 2 (ERR2), endoplasmic reticulum processes (GORS1, MA1B1), cellular death and repair (DAPK2, RD23A), with downregulated phosphodiesterase activity (PDE3A, PDE7A), calcium/calmodulin genes (CAC1I, PDE1C, PLPL9, RAMP1), adenylate cyclase activity (ADCY2,7,9), lipids and fatty acid processes (ACBG2, DGKD, DGKZ). Furthermore, melanin hormone activity (MCH, MCH2) was the only hormone-related process detected in this condition and exclusively in HB. Finally, associative learning was the only cognitive process found in the combined condition, underlined by immediate early genes (FOS, FOSL2), synapse-regulation genes (NLGNX, NPTX1), long-term potentiation plasticity (NPTX2), osmoregulation and corticotropin releasing factor regulation (UTS1), but dopamine nor serotonin were not differentially expressed in this condition. Unlike Warming and High CO_2_ in isolation, no changes in behaviour or memory-related mechanisms were exhibited in this combined condition.

#### b) Acanthurus leucosternon

Client individuals exposed to Warming & High CO_2_ displayed a relatively small molecular response compared to the cleaner wrasse (figure 3b & 5). The molecular reprogramming was though mostly shared (72%) with Warming. This combined condition elicited unique differential expression of microtubule nucleation, binding, bundle activity, and as well as organ and tissue development. Moreover, locomotory behaviour and medium-term memory were found in the client in contrast with the cleaner wrasse, while associative learning was observed in both species. There were several responses to stress, including positive regulation of response to endoplasmic reticulum stress and apoptosis, but no cellular death functions were exhibited. In particular, molecular mechanisms of cell growth, glutamatergic synapses, immune response, immediate early genes, apoptosis, and DNA damage were part of the transcriptional processes altered under this condition in the FB and HB (table S11). Even though no enriched functions were found significant in MB, the differential gene expression found was related to DNA binding, transcription factor, and GABA (table S11).

## DISCUSSION

Future environmental conditions of Warming and High CO_2_ elicit molecular reprogramming in the brains of the interacting cleaner wrasse *L. dimidiatus* and its client *A. leucosternon.* Here, the cleaner wrasse *L. dimidiatus,* in particular, exhibited large transcriptional alterations and a lower interaction motivation and quality. A disruption to the mutualistic cleaning behaviour in future environmental conditions has also been previously observed (Paula et al., 2019b). Moreover, the loss of cleaners’ cognitive and strategic sophistication has been associated with increasing CO_2_ levels (J. Paula, Baptista, et al., 2019) and back-to-back extreme environmental disturbances (cyclones, heatwaves, and bleaching; Triki et al., 2018). This combined evidence shows that future environmental conditions will likely disturb this important mutualistic cleaning behaviour. One of the reasons for stressor-led behavioural change may be the large neuromolecular alterations displayed, especially for *L. dimidiatus*, when exposed to future conditions. This hints at a need for more adjustments in physiological functions and an elevated impact of environmental changes on species with elevated cognitive abilities involved in their survival. Here we shed light on the expression patterns underlying the cleaning interaction in predicted near-future environmental conditions and reveal transcriptional and functional mechanisms underlying the susceptibility of this vital cleaning mutualism.

Warming revealed the largest transcription of cellular stress responses for both species involved in the mutualistic interaction. Cellular stress activity is commonly transcribed with repair mechanisms (*e.g.* immune response, ATP-related processes; Morimoto, 1993). DNA damage and apoptosis initiation genes were expressed at elevated levels along with heat shock proteins, which are important in maintaining cellular activity and preventing DNA degradation (Logan & Buckley, 2015). Such processes are costly, and alterations in metabolic actions are expected (Liu et al., 2016). For instance, significant reductions of energy metabolites (*e.g.* DOPAC and 5-HIAA) have been previously found underlying *L. dimidiatus* motivation and quality to interact (Paula et al., 2019b). Expression changes in glycine and glutamate during Warming emphasize osmotic and energetic disruptions that may limit the transportation of glucose and oxygen from the blood to the brain (Schmidt et al., 2017). Genes related to hypoxia-inducible factors and further osmolytes, such as lactate, were highly upregulated uniquely under Warming in our cleaner wrasse, suggesting high-stress levels and an alteration of osmolytes and metabolites. This may have the potential to compromise behavioural performance, as seen in reduced activity levels, cognitive performance, muscular activity, and lateralization behaviour in other fishes with elevated temperature (Biro et al., 2010; Jarrold & Munday, 2018; Maille & Schradin, 2017; Pörtner & Knust, 2006; Schmidt et al., 2017). The behavioural changes in the cleaning interaction during Warming, however, may not be a direct cause of elevated temperature interfering with the capacity to interact in the brain, but more indirect by needing to shift focus on increased metabolism related to cellular stress responses and oxygen demands in the brain (Mattiasen et al., 2020; Song et al., 2019).

High CO_2_ altered responses of stimulus reception, ion transport, circadian rhythm, visual perception, dopamine, and neurotransmitters (glutamate and GABA) in the brain of *L. dimidiatus*, indicating an effect of acidification on neurotransmission. While GABA receptors were differentially expressed in all three conditions, only High CO_2_ showed an effect on most of the pathway, including the upregulation of genes associated with GABA signalling, sodium and chloride-dependent transporter and GABA aminotransferase (GABT). The upregulation of GABT facilitates the degradation of GABA into succinic semialdehyde, which is in charge of regulating the supply of GABA in the brain (“GABA shunt”; Deutch & Roth, 2004). As a result, succinic semialdehyde is incorporated into the Krebs cycle from which glutamine is formed and subsequently converted into glutamate (Olsen & Li, 2012), a major excitatory neurotransmitter already known to be regulating the interaction behaviour in the cleaner wrasse (Ramírez-Calero et al., 2021). Our results thus suggest an interplay between the expression of GABAergic and glutamatergic pathways in the cleaner wrasse when faced with ocean acidification. GABA ion channel interference in High CO_2_ has been attributed to neurotransmission dysfunction (Schunter et al., 2018, 2019) in the brain of teleost fish. In fact, administration of a GABA_A_ receptor antagonist (gabazine) led to the recovery of most High CO_2_-induced cleaning behaviour alterations (Paula et al., 2023). Moreover, under control conditions, the administration of a GABA_A_ receptor agonist (muscimol), produced similar cleaning motivational disruptions as High CO_2_, suggesting a crucial role of the GABA ion channel interference as a mechanism for cleaning behaviour disruption under High CO_2_. A variety of alterations in lateralization behaviour (Nilsson et al., 2012), predator recognition and learning performance (Chivers et al., 2014) are also known to be related to GABAergic neuromodulation. Another important neurotransmitter in the brain, dopamine, is also connected to the interaction behaviour in *L. dimidiatus* and plays a key role already in normal environmental conditions (Ramírez-Calero et al., 2021). Alterations (significant reductions) of dopamine activity have been observed under High CO_2_ and connected to a reduction in motivation to interact as well as the reward and risk perception system (de Abreu et al., 2020; Paula et al., 2019b; Soares et al., 2017b). The gene expression changes of dopamine receptors under High CO_2_ suggest transcriptional modifications that are being altered in the dopaminergic pathway, which is vital for *L. dimidiatus* success in consolidating cleaning behaviour. Thus, changes in glutamatergic/GABAergic and dopaminergic neurotransmission and ion transport under acidification could explain the disruption of the mutualistic interaction with ocean acidification.

Alterations in GABA neurotransmission and ion transport have also been associated with visual functions (Chung et al., 2014; Nilsson et al., 2012). Changes in ion gradients over neuronal membranes through GABA receptor activity under High CO_2_ were linked with reduced retinal reaction time (and reduced speed in light response) in another coral reef fish (*Acanthochromis polyacanthus*). *L. dimidiatus* upregulated genes involved in visual perception, including key photoreceptors, retinal development genes, and GABA receptors, under High CO_2_. These changes may suggest an alteration in retinal reaction time due to a disruption of retinal action potentials (*e.g.* V-ATPase pump; Damsgaard et al., 2020; Hirasawa et al., 2012), as fish vision is regulated by changes in the intra and extra-cellular pH of the retina modulated by the vacuolar adenosine triphosphatase and the expression of photoreceptors (V-ATPase; Damsgaard et al., 2020). Consequently, we hypothesize that this transcriptional modification may alter stimuli reception in ganglion cells and the retina reducing its normal reaction time (Chung et al., 2014). This response may also be related to changes in the circadian rhythm found altered for *L. dimidiatus*. High CO_2_ commonly alters the transcriptional regulation of the circadian pathway in fish brains across many species, including coral reef fishes (Kang et al., 2022; Schunter et al., 2016, 2021; Suresh et al., 2023). Although further studies are needed to corroborate the link and its mechanism under ocean acidification, pH changes have the potential to alter the visual perception of cleaners, including the speed of processing images, which is a crucial ability to recognize clients as well as consolidate long-standing cleaning relationships (Tebbich et al., 2002).

While it is important to understand the mechanisms underlying the responses to near-future conditions in isolation, revealing antagonism and synergism of multiple drivers is crucial for identifying molecular processes in more realistic future environmental conditions (Munday et al., 2019; Nagelkerken & Munday, 2016). We found that much of the transcriptional changes were shared with Warming, including metabolic genes and immune response, whereas ion regulation, GABA/glutamatergic activity and synaptic transmission were processes shared with High CO_2_. Despite sharing a transcriptional response with the single treatments, the combination of the future conditions revealed enhanced stress signatures underlined by endoplasmic reticulum (ER) stress response in the cleaner wrasse, suggesting the transcription of protein folding regulation and homeostasis. This type of stress can alter synaptic function and memory storage due to exposure to harmful stimuli, such as hypoxia and oxidative stress caused by environmental stress (Díaz-Hung et al., 2020). Cellular stress responses have been observed in other fish, suggesting increasing mortality and failure of acclimation through phenotypic plasticity (Feidantsis et al., 2015; Shi et al., 2019). Thus, the elevated stress signatures observed in the cleaner wrasse show increased demands of protein folding and homeostasis repair, signalling, synaptic function, and regulation of organ development imposed by both combined environmental changes. As these modifications are not detected in conditions in isolation, it seems this may pose an additive effect in response to future environmental conditions and has the potential to disrupt important biological functions.

Interestingly, while signs of stress increased in the combined condition, there were no expression changes in behaviour-related genes. The proportion of time interacting did not significantly decrease during Warming & High CO_2_ compared to control. This observation suggests that fish may prioritize maintaining biological performance (e.g. protein folding and repair) while making fewer adjustments to the transcriptional response underlying the interaction behaviour. Trade-offs between biological performance and behaviour have been documented in other fish species revealing limits in the adaptive potential to climate change (Allan et al., 2017; Domenici et al., 2014; Laubenstein et al., 2019; Paula et al., 2019b). Combined environmental stressors have been shown to modify correlations between physiological function and behavioural traits through differential phenotype sensitivities and environmental contexts (Killen et al., 2013). In the cleaner wrasse, molecular mechanisms underlined by increased ER stress, organ development, and morphogenesis compared to reduced glutamatergic neurotransmission and an absence of changes in behavioural genes and dopamine suggest a trade-off between these two processes (i.e. stress and behaviour). As such, the trade-offs between physiological performance related to stress and behaviour in the context of mutualistic interactions could compromise the establishment of future social interactions in *L. dimidiatu*s, and its ability to adapt to climate change.

In conclusion, the molecular signatures underlying the mutualistic interaction of *L. dimidiatus* and a client species *A. leucosternon* under predicted future climate change conditions provoked an array of effects that compromised cleaning interactions, ranging from cellular stress responses, alteration of neurotransmission, and high metabolic demands. During Warming, the upregulation of hypoxia, osmolyte, and metabolite alterations suggested homeostatic disruption and aerobic constraints due to high temperatures. Furthermore, exposure to High CO_2_ altered gene expression in ion transport and neurotransmission driven by GABAergic and glutamatergic neurotransmission. This suggests that an interplay between these two processes also affects motivation to interact in *L. dimidiatus*. In addition, under High CO_2_, visual perception genes were altered in expression, potentially disrupting social relationships as cleaner wrasses strongly rely on client recognition. Finally, the more realistic future combined condition of elevated Warming & High CO_2_ created ER and apoptotic stress on the one hand but also the absence of molecular signatures related to behaviour and less behavioural impairments on the other hand, suggesting a trade-off between these physiological functions and behaviour in *L. dimidiatus.* Altogether the molecular and behavioural responses to climate change-related future environments can compromise the establishment of future social interactions in the cleaner wrasse and its ability to adapt to increased ocean acidification and warming. Overall, this can have major implications in the future for cleaning interactions and the key ecosystem services they provide.

## MATERIALS AND METHODS

### Experimental setup

To identify the functional molecular basis of the interaction between two fish species under ocean acidification and Warming conditions, 24 female adult individuals of *L. dimidiatus* and 24 females of *A. leucosternon* were collected by TMC-Iberia in the Maldives islands and transported to the aquatic facilities of Laboratório Marítimo da Guia (MARE) in Cascais, Portugal. We selected the fish species *A. leucosternon* as a client since it is one of the most frequent clients for the genus *Labroides* (Tebbich et al., 2002). For *L. dimidiatus*, female individuals (∼7 cm) were used to reduce the effect of sex as this species is protogynous (Kuwamura et al., 2002). Furthermore, fish were deparasitized with a freshwater bath upon arrival, cleaner wrasses were kept separately in individual tanks (20 L) to avoid aggressive behaviours and surgeonfish (*A. leucosternon*) were kept in groups. All fish were fed *ad libitum* once per day, and further information on size and weight can be found in table S1. Each individual was first laboratory-acclimated for five days in seawater conditions similar to their native site (salinity = 35 ± 0.5), temperature 29 °C (Maldives 2013–2014 average SST, NOAA), pH 8.1 and pCO_2_ 400 ppm (2014 BOBOA Ocean Acidification mooring, NOAA). Following acclimation, each fish was exposed to one of four experimental treatments for 28 days, namely: (1) present-day scenario (control) (29 °C, pH 8.1, pCO_2_ ∼400 ppm), (2) Warming (32 °C, pH 8.1, pCO_2_ ∼400 ppm), (3) High CO_2_ (29 °C, pH 7.7, pCO_2_ ∼1000 ppm) and (4) Warming & High CO_2_ (32 °C, pH 7.7, pCO_2_ ∼1000 ppm), following IPCC’s RCP scenario 8.5 (Bindoff et al., 2019, figure 1, table S1-S2)

Experimental tanks had a semi-open flow-through aquatic system to maintain alkalinity levels, dissolved carbon and pH. Natural seawater was UV-irradiated with a Vecton V2 300 Sterilizer before being passed to each experimental tank. Photoperiods of 12 h/12 h with light and dark cycles were maintained. Ammonia and nitrate levels were checked daily using colourimetric tests (Salifert Profi Test, Holland), pH levels were automatically monitored and adjusted every two seconds (Profilux 3.1 N, GLH, Germany), regulated by injection and aeration of certified CO_2_ gas and filtered atmospheric air, respectively (Air Liquide, Portugal; soda lime, Sigma-Aldrich). Seawater temperature was regulated using underwater heaters 300 W, TMC-Iberia (Portugal). Additional equipment was used to complement the daily monitoring of seawater temperature (VWR pH 1100H pH meter, Avantor, US), salinity (V2 refractometer TMC Iberia, Portugal) and pH (826 mobile pH meter, Metrohm, Germany). Additional quantifications of pH were done using a pH meter connected to a glass electrode (Schott IoLine, Si analytics, ±0.001), calibrated with TRIS-HCl (TRIS) and 2-aminopyridine-HCl (AMP) seawater buffers. Finally, Seawater carbonate system speciation was calculated twice a week from total alkalinity (spectrophotometrically at 595 nm) and pH measurements. Bicarbonate and pCO_2_ values were calculated using CO2SYS software (table S1).

### Behavioural analysis

Following the 28-day period of exposure to the treatment conditions, we initiated the behavioural tests. These tests were conducted in specially set up observation tanks (40 L) situated in a designated observation room. Before the interaction, both cleaners and clients underwent a 24-hour fasting period. The behavioural trial entailed placing one cleaner wrasse and one client into an observation tank that replicated the environmental conditions of their respective treatments. Each trial was recorded for a duration of 40 minutes, discounting the initial 5 minutes allocated for acclimation. The analysis of the behavioural trials was carried out following Paula et al., (2019b). Cleaning behaviour was divided into two primary components: (i) motivation to interact and (ii) interaction quality. To characterise motivation to interact, we measured the proportion of time spent in interaction (close body inspection and removal of damaged tissue or scales), the proportion of interactions initiated by the cleaners, and the ratio of the client “posing” displays (*i.e.* client “posing” displays/time of no interaction; “posing” displays are conspicuous signals used by clients seeking cleaning interactions).

We evaluated the quality of the interactions based on the duration of the interaction, the frequency of client jolts per 100 seconds of interaction (these jolts are notable signals that imply cheating or dishonesty on the part of the cleaner), and the proportion of interaction time that was spent on tactile stimulation events (touches with pectoral fins known to alleviate stress and prolong interaction duration). We used the event-logging software “Boris” to analyse all behavioural videos, with the behavioural catalogue detailed in supplementary table S3a as a reference. Both the cleaner fish and the client were considered as focal subjects in the analysis (Friard & Gamba, 2016). Following the behavioural trial, each fish was euthanized. We then measured the body length and extracted the three main regions of the brain for further investigation. The extracted tissues were stored at -80°C for subsequent processing.

### Statistical analysis of behavioural data

Data exploration was performed according to Zuur et al., (2010), which promotes a protocol for data exploration. We analysed behavioural data using generalised linear models (GLM) with a Gaussian distribution. These models used CO_2_ treatment (factor with two levels: Control; High CO_2_) and temperature treatment (factor with two levels: 29°C and 32°C) as categorical fixed factors, according to Zuur & Ieno (2016) The full models, with all possible interactions, were tested using the function “glm” and the function “Anova” from the package “car” (Fox & Weisberg, 2019) in R, version 3.4.3 (R Core Team, 2017). *Post-hoc* multiple comparisons were performed using the package “emmeans” (Lenth, 2020) with Benjamini-Hochberg corrections. Model assumptions and performance were validated using the package “performance” (Lüdecke et al., 2021). Data exploration used the HighstatLibV10 R library from Highland Statistics (Zuur et al., 2009).

### RNA extraction and sequence processing

Total RNA was extracted using the RNAeasy Mini Kit protocol (Qiagen). Homogenization with sterile silicon beads was performed at maximum speed for 30 seconds in a Tissuelyzer (Qiagen), and Dnase I treatment (Qiagen) was performed according to the manufacturer’s protocol to remove DNA contaminants. The quantity and quality of RNA were tested using a Qubit fluorometer and an Agilent Bioanalyzer for RNA Integrity Numbers (RIN). Samples with a RIN > 8 were retained only. mRNA-focused sequencing libraries were designed with Illumina TruSeq v3 kits and sequenced for paired-end reads of 150 bp length on an Ilumina Hiseq4000 at the King Abdullah University of Science and Technology corelab facility.

To assess the differential gene expression of species interactions under different environmental conditions, raw read quality was examined using FastQC v. 0.11.9 (Andrews, 2010). Further, poor-quality sequences were trimmed, and adapters were removed using the software Trimmomatic v.0.36 (ILLUMINACLIP:TruSeq3-PE.fa:2:30:10 LEADING:4 TRAILING:3 SLIDINGWINDOW:4:15 MINLEN:40; Bolger et al., 2014). A *de novo* transcriptome assembly from a previous study (Ramírez-Calero et al., 2021) was used for each species separately, and the functional annotation of transcripts was conducted using BLAST+ 2.10.0, Swissprot/Uniprot protein database (November 29, 2019) and Zebrafish (*Danio rerio,* Apr 2018). In addition, the Ballan wrasse genome annotation (*Labrus bergylta*, March 2020, GCA_900080235.1) was used for *L. dimidiatus* assembly only as it is the closest species to the cleaner wrasse with a reference available. Omicsbox v. 1.3 (Götz et al., 2008) was used to functionally annotate the transcripts with Gene Ontology (GO terms) and KEGG pathways.

### Differential Expression Analyses

Quantification of transcript abundance for each species was obtained using the script *align_and_estimate_abundance.pl* from Trinity software. RSEM v1.3.3 (Li & Dewey, 2011) was set as quantification method and Bowtie2 as mapping tool (Langmead & Salzberg, 2012) using *–gene_trans_map* to receive gene-level counts. Gene expression matrices for both species were built with the script *abundance_estimates_to_matrix.pl* (Haas et al., 2013), while low expression transcripts (<10 read counts) were filtered out with the *filter_low_expr_transcripts.pl* script. The command *–highest_iso_only* was used to retain the most highly expressed isoforms for each gene.

To statistically evaluate differential gene expression, we used DESeq2-package v. 1.26.0 (Love et al., 2014) in R with a Wald test statistic using the model *design = ∼brain_region+treatment* to evaluate the expression differences for each treatment (Control; High CO_2_; Warming; Warming & High CO_2_) factoring in the different brain regions (Forebrain; Midbrain; Hindbrain). As brain regions show large differences in gene expression, further models were used (*design =* ∼*treatment*) where each brain region was examined separately. Resulting outliers were examined using Principal Component Analysis (PCA), and two individuals of *A. leucosternon* from the control condition were removed (figure S7-S8). For each species, pair-wise comparisons were computed by comparing environmental conditions to the control: control vs Warming, control vs High CO_2_, and control vs Warming & High CO_2_. We accepted a gene to be differentially expressed with an FDR p-adjusted value of 0.05 and an absolute log2fold change threshold of 0.3. Functional enrichment was performed using Fisher’s exact test by testing the resulting subsets of differentially expressed genes against all transcripts in the transcriptome using an FDR significance value of 0.05 in Omicsbox v. 1.3.

## DATA AVAILABILITY

The data that support the findings of this study containing the raw sequencing files and *de novo* transcriptome assemblies (*Labroides dimidiatus* and *Acanthurus leucosternon*) are openly available under the NCBI BioProject PRJNA72634 and accession number GJED00000000.

## ETHICS STATEMENT

Research was conducted under approval of Faculdade de Ciências da Universidade de Lisboa animal welfare body (ORBEA – Statement 01/2017) and Direção-Geral de Alimentação e Veterinária (DGAV – Permit 2018-05-23-010275) in accordance with the requirements imposed by the Directive 2010/63/EU of the European Parliament and of the Council of 22 September 2010 on the protection of animals used for scientific purposes.

## Acknowledgements

We would like to thank Celia Schunter’s lab members at HKU Sneha Suresh, Jingliang Kang, Natalia Petit-Marty, Arthur Chung and Jade M Sourisse for their help in improving the bioinformatic pipeline and data analysis, and for engaging in stimulating discussions and comments to this work. We also would like to acknowledge Lígia Cascalheira and Dr Tiago Repolho for their help with the maintenance of aquatic systems throughout the experiment.

This work was supported by the Hong Kong Research Grant Committee Early Career Scheme fund 27107919 (CS) and FCT—Fundação para a Ciência e Tecnologia, I.P., within the project grant PTDC/BIA-BMA/0080/2021 – ChangingMoods, the scientific employment stimulus program 2021.01030. CEECIND, the strategic project UIDB/04292/2020 granted to MARE, project LA/P/0069/2020 granted to the Associate Laboratory ARNET, and the PhD scholarship UI/BD/151019/2021 awarded to EO.

