## Supplementary Figures for "Neuromolecular responses in disrupted mutualistic cleaning interactions under future environmental conditions"

Running title (45 characters): Cleaning interactions in climate change.

**SUPPLEMENTARY FIGURES**

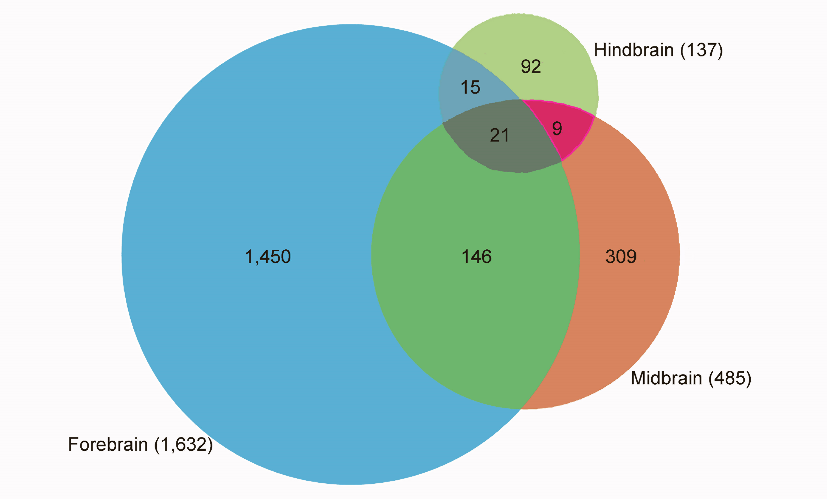

**Figure S1.** Unique and overlapping differentially expressed genes (DEGs)present in the Warming condition for *L. dimidiatus*

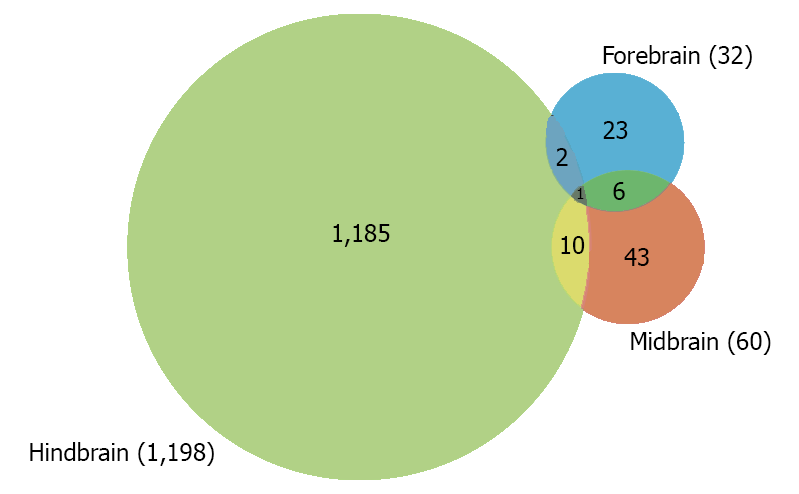

**Figure S2.** Unique and overlapping differentially expressed genes (DEGs) present in the High CO_2_ condition for *L. dimidiatus*

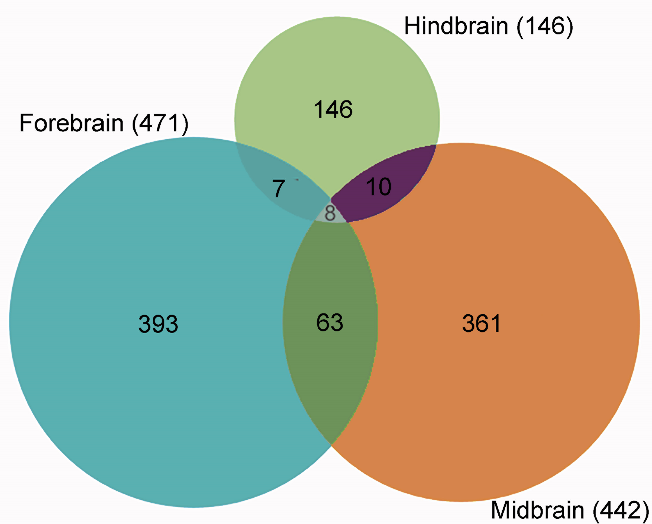

**Figure S3**. Unique and overlapping differentially expressed genes (DEGs) present in the Warming & High CO_2_ condition for *L. dimidiatus*.

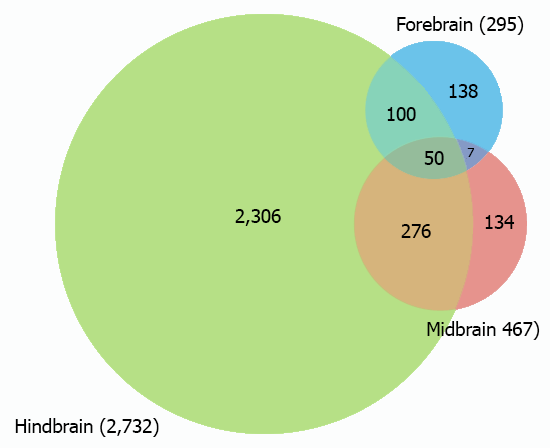

**Figure S4.** Unique and overlapping differentially expressed genes (DEGs) present in the Warming condition for *A. leucosternon.*

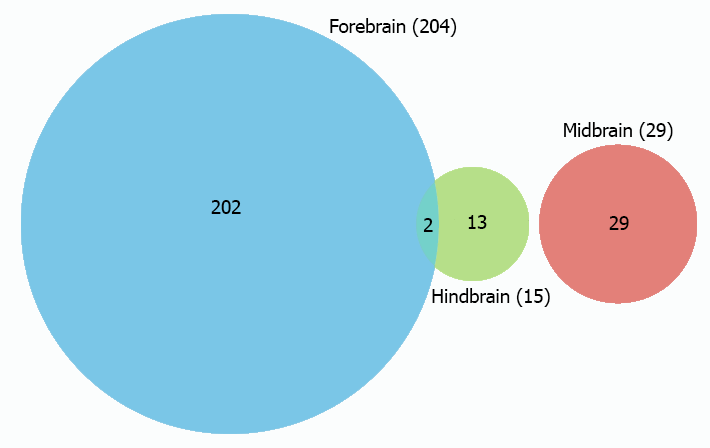

**Figure S5.** Unique and overlapping differentially expressed genes (DEGs) present in the High CO_2_ condition for *A. leucosternon.*

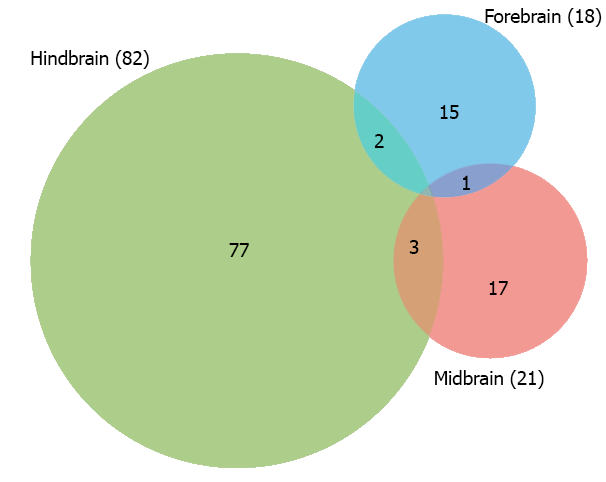

**Figure S6.** Unique and overlapping differentially expressed genes (DEGs) present in Warming & High CO_2_ condition for *A. leucosternon.*

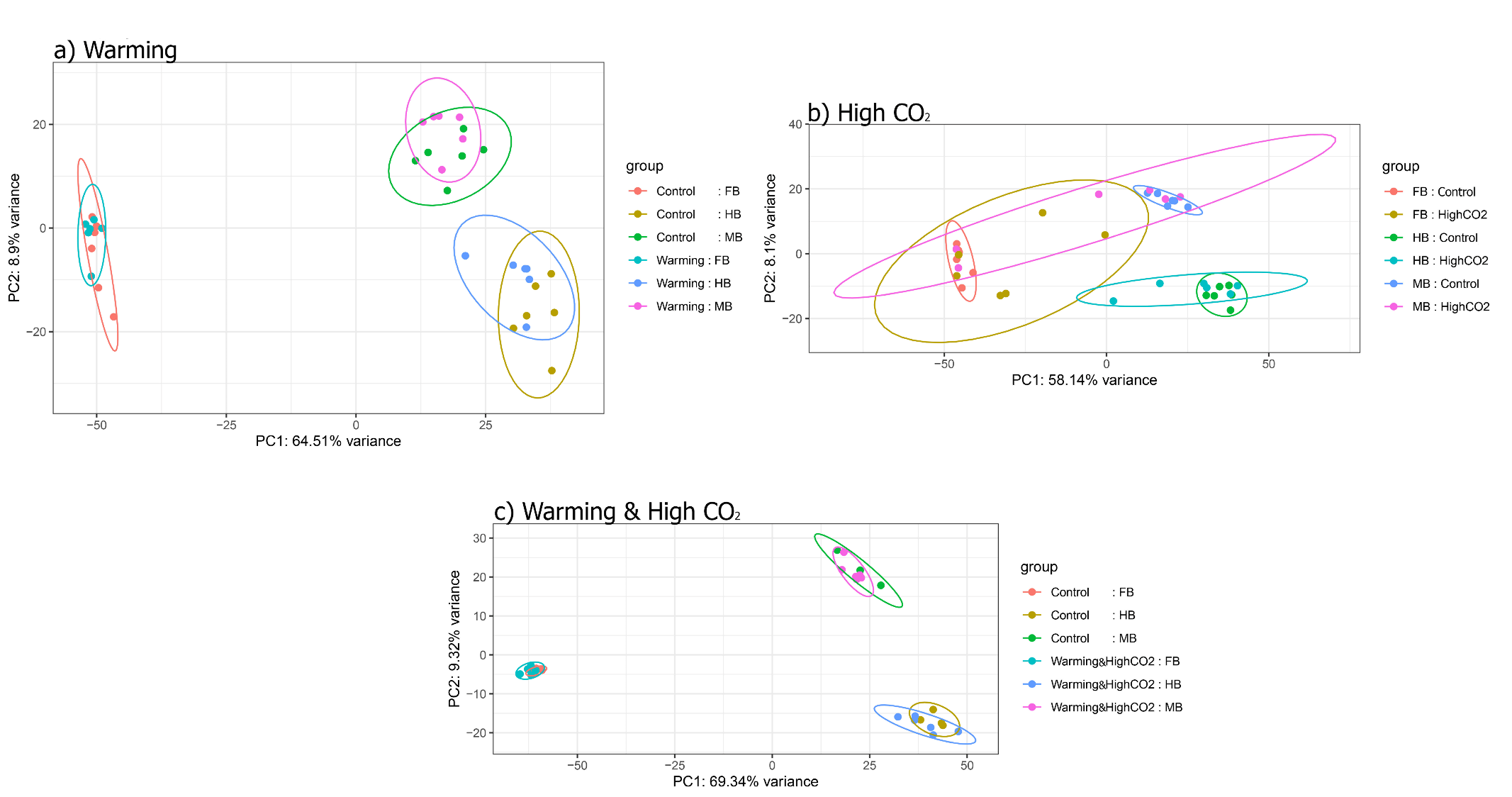

**Figure S7**. PCA’s of normalized gene counts using the design *~brain_region + treatment* and a *rlog* transformation (rld) for each environmental treatment: a) Warming, b) High CO_2_ and c) Warming & High CO_2_) for *A. leucosternon.*

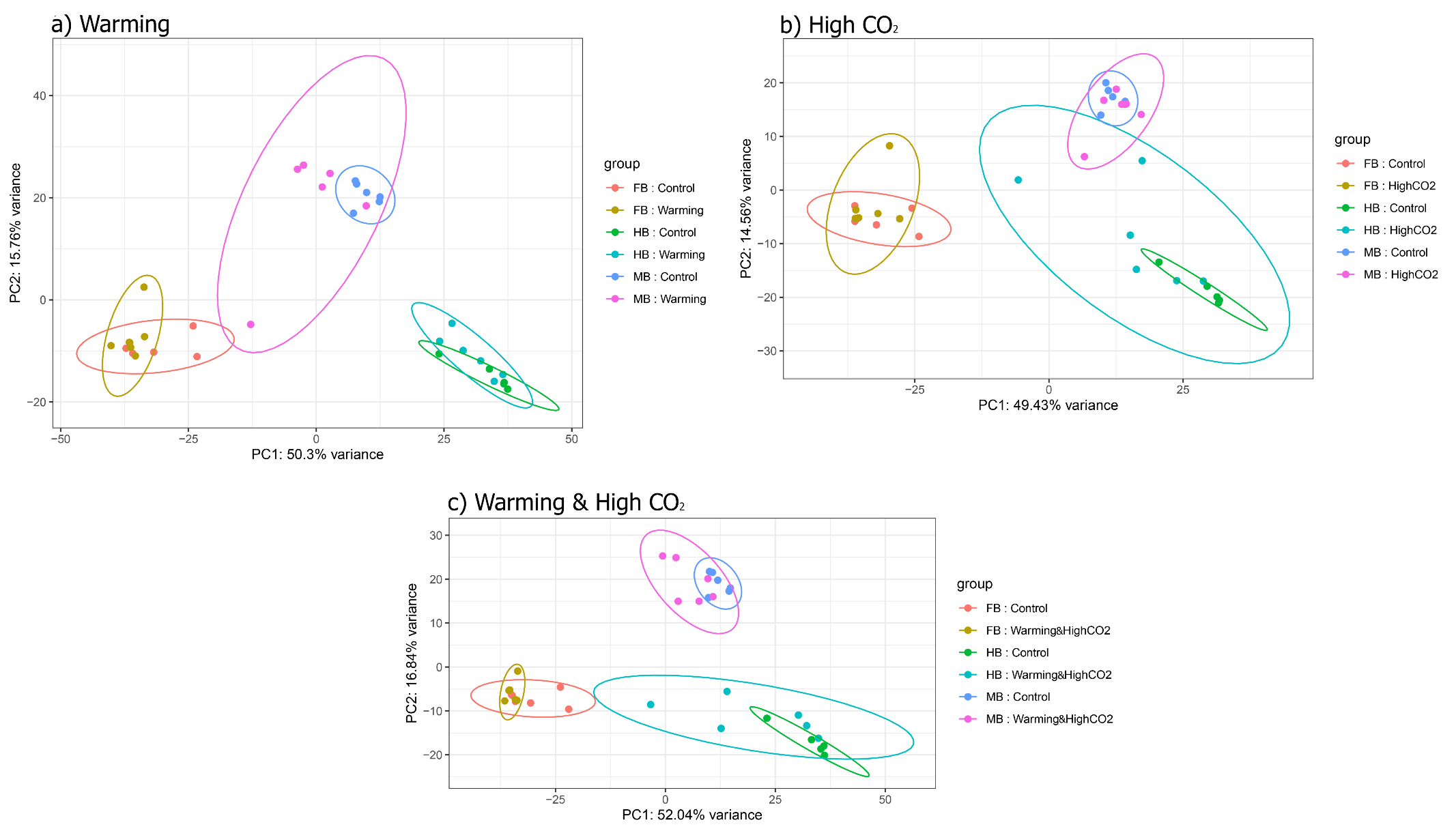

**Figure S8**. PCA’s of normalized gene counts using the design *~brain_region + treatment* and a *rlog* transformation (rld) for each environmental treatment: a) Warming, b) High CO_2_ and c) Warming & High CO_2_ for *L, dimidiatus*.
